# Looking at my own Face: Visual Processing Strategies in Physical Self-representation

**DOI:** 10.1101/096073

**Authors:** Anya Chakraborty, Bhismadev Chakrabarti

## Abstract

We live in an age of ‘selfies’. Yet, how we look at our own faces has seldom been systematically investigated. In this study we test if visual processing of self-faces is different from other faces, using psychophysics and eye-tracking. Specifically, the association between the psychophysical properties of self-face representation and visual processing strategies involved in self-face recognition was tested. Thirty-three adults performed a self-face recognition task from a series of self-other face morphs with simultaneous eye-tracking. Participants were found to look at lower part of the face for longer duration for self-face compared to other-face. Participants with a reduced overlap between self and other face representations, as indexed by a steeper slope of the psychometric response curve for self-face recognition, spent a greater proportion of time looking at the upper regions of faces identified as self. Additionally, the association of autism-related traits with self-face processing metrics was tested, since autism has previously been associated with atypical self-processing, particularly in the psychological domain. Autistic traits were associated with reduced looking time to both self and other faces. However, no self-face specific association was noted with autistic traits, suggesting that autism-related features may be related to self-processing in a domain specific manner.

## Introduction

Self-awareness is one of the most complex manifestations of human cognition and argued to be a prerequisite for understanding mental states of ‘self’ and ‘others’(Gallup 1970, Keenan, McCutcheon et al. 1999).Self-awareness exists in different domains, e.g. in the physical domain as the awareness of one’s own body and face, in the psychological domain as an entity with specific traits and qualities, and in the temporal domain as a continuous being across time(James 1890). Physical self-awareness is one of the earliest and most basic domains of self-awareness to develop. Among other methods, self-face recognition has been often used as a paradigm to interrogate this domain of self-processing(Amsterdam 1972, Keenan, Ganis et al. 2000, Sugiura, Kawashima et al. 2000, Kircher, Senior et al. 2001, Keenan, Wheeler et al. 2003, Nielsen and Dissanayake 2004, Uddin, Kaplan et al. 2005, Breédart, Delchambre et al. 2006, Sui, Zhu et al. 2006, Platek, Wathne et al. 2008, Devue, Van der Stigchel et al. 2009, Pannese and Hirsch 2011) and most studies on physical self-representation have focussed on the investigation of the behavioural and neural basis of self-face recognition(Keenan, McCutcheon et al. 1999, Tong and Nakayama 1999, Kircher, Brammer et al. 2002, Uddin, Kaplan et al. 2005, Platek, Wathne et al. 2008, Ma and Han 2010).Comparatively little is known(Kita, Gunji et al. 2010, Hungr and Hunt 2012) about gaze behaviour during the recognition of a face as belonging to oneself, leading to the question of whether the gaze response pattern for a face recognized as ‘self’ is different from one recognized as ‘other’. Are self-faces special or merely familiar faces, as would be argued by some theoretical accounts(Gillihan and Farah 2005)? The study of eye gaze behaviour in self-face recognition allows for better understanding of visual strategies underpinning this fundamental aspect of physical self-awareness. In an age of selfies, how we look at our own face assumes an importance beyond the academic domain.

Looking at self-face is associated with greater attention to and faster recall compared to other faces (Tong and Nakayama 1999, Devue, Van der Stigchel et al. 2009). Identification of self-face requires orientation towards the self from a decentralized position and indicates high salience for self-related stimuli(Heinisch, Dinse et al. 2011). The self-face is identified faster among other faces even where faces are presented in non-upright conditions (Tong and Nakayama 1999) (though see Devue et al(Devue, Van der Stigchel et al. 2009)).Such high salience for self-related stimuli is also evident from their facilitatory effect on spatial priming(Pannese and Hirsch 2011), interference with cognitive tasks(Breédart, Delchambre et al. 2006), as well as from EEG experiments showing an increased P300 signal (related to attention allocation) to self-name (Gray, Ambady et al. 2004). Self-specific stimuli have been found to alter the salience of neutral stimuli by association leading to the proposal of the Self-Attention Network (SAN)(Sui and Humphreys 2013, Porciello, Minio-Paluello et al. 2016). SAN constitute a model where neural networks involved in self-processing interact with attentional networks to determine self-salient behaviour. Notably, this individual-specific salience for the self-face is distinct from the salience associated with low-level visual features of the presented stimulus.

Different gaze allocation strategies are employed to discriminate between familiar and unfamiliar faces. Notably, these strategies are task-dependent, e.g. when viewing unfamiliar faces in a recall task, participants showed greater sampling from all regions of the face while greater gaze fixation to the eye region was noted for more familiar faces(Heisz and Shore 2008). In contrast, participants showed increased sampling and exploration of facial features for familiar faces compared to unfamiliar faces during a recognition task(Van Belle, Ramon et al. 2010). In order to investigate if self-faces are visually processed similar to familiar faces, the current study uses a self-other face recognition task to investigate gaze response pattern for faces identified as belonging to self compared to those identified as ‘other’. It is predicted that if self-faces are processed similarly to other familiar faces then viewing the self-face should be associated with increased sampling and exploration of facial features of the self-face.

Self-face representation has behaviourally been studied using psychophysics paradigms that utilise morphed stimuli and calculate a psychometric response function based on the participant’s identification of the morphs as ‘self’ or ‘other’(Chakraborty and Chakrabarti 2015). However, such paradigms tend not to include simultaneous eye-tracking measures; it is thus not possible to elucidate if the pattern of gaze fixation to self and other faces can predict the mental representation of the self-face. Using a morphing paradigm, an eye-tracking study reported a stronger gaze cueing effect for self-similar compared to self-dissimilar faces(Hungr and Hunt 2012). However, visual processing strategies of self and unfamiliar faces in previous studies have not been mapped onto the psychometric properties of self-other face representation. The current study addresses this gap in knowledge by using a self-other face morphing paradigm to investigate self-face representation with simultaneous eye-tracking.

A secondary aim of the current study tested individual differences in self-face processing in relation to autism-related traits. Atypical gaze fixation to social stimuli(Klin, Jones et al. 2002, Pelphrey, Sasson et al. 2002, Dalton, Nacewicz et al. 2005) as well as atypical self-processing (Lombardo, Barnes et al. 2007, Uddin, Davies et al. 2008, Lombardo, Chakrabarti et al. 2009, Lombardo, Chakrabarti et al. 2010)is well documented in individuals with Autism Spectrum Disorders (ASD). Accordingly, the current study aimed to test if there was any association between autistic traits and gaze duration to faces in general and if any such association is specific to facial identity (self or other) and/or facial region (upper vs. lower parts of the face). Measurement of autistic traits in the general population can help investigate how autistic symptoms map onto social behaviour. Autistic traits are distributed continuously across the population, and individuals with ASD score highly on these measures (Baron-Cohen, Wheelwright et al. 2001). Individuals with a clinical diagnosis of ASD typically score at the high end of this continuous distribution of autistic traits. Measuring autistic traits in the general population allows one to measure associations between trait features and experimental manipulations, thus providing an initial foundation for follow-up investigations with the clinically diagnosed tail of the trait distribution (Robinson, Koenen, McCormick, et al., 2011). In the present study, autistic traits are measured using Autism Spectrum Quotient (AQ) (Baron-Cohen, Wheelwright et al. 2001). AQ scores range from 0 to 50, and individuals scoring 32 or higher have >80% chance of having an ASD diagnosis (Baron-Cohen, Wheelwright et al. 2001). AQ has been developed based on the behavioural symptoms of ASD, and has five subdomains that include social skills, attention switching, attention to detail, communication, and imagination.

In view of the evidence discussed above for different gaze fixation patterns for familiar and unfamiliar faces, the first question of interest is whether gaze duration for upper parts of the face (including eye region) and for lower parts of the face (including mouth) differs between faces identified as the self vs. those identified as other? Following the reports of visual processing of familiar faces, it is predicted that gaze duration to the lower parts of the face will be longer for a face identified as ‘self’ compared to ‘other’.

The second aim is to investigate the relationship between self-face representation at the behavioural level (operationalised by the slope of the psychometric function) and gaze metrics for faces identified as ‘self’ versus ‘other’. In the context of the present study, the ‘self’ and ‘other’ constitute two categories. The steepness of the slope of the psychometric function for the self-face recognition curve (derived from self-other face morphs) provides a measure of the overlap between the two categories. A steeper slope of the self-face recognition curve indicates a lower extent of overlap between self and other categories, i.e. amore distinct self-face representation. In other words, it would take small changes in stimulus feature (changes in morph percentages) to shift from the self to other category for an individual with a more distinct self-face representation. Conversely, a shallower slope of the self-face recognition curve indicates a broader spread of category boundary that requires larger change in stimulus feature to shift from the self to other category (Fig. 1). The ability to recognise oneself physically, specifically through self-face is believed to be associated with prosocial behaviours like empathising and emotion recognition that require both self-other overlap and self-other distinction (Bird and Viding 2015).

**Fig. 1.**
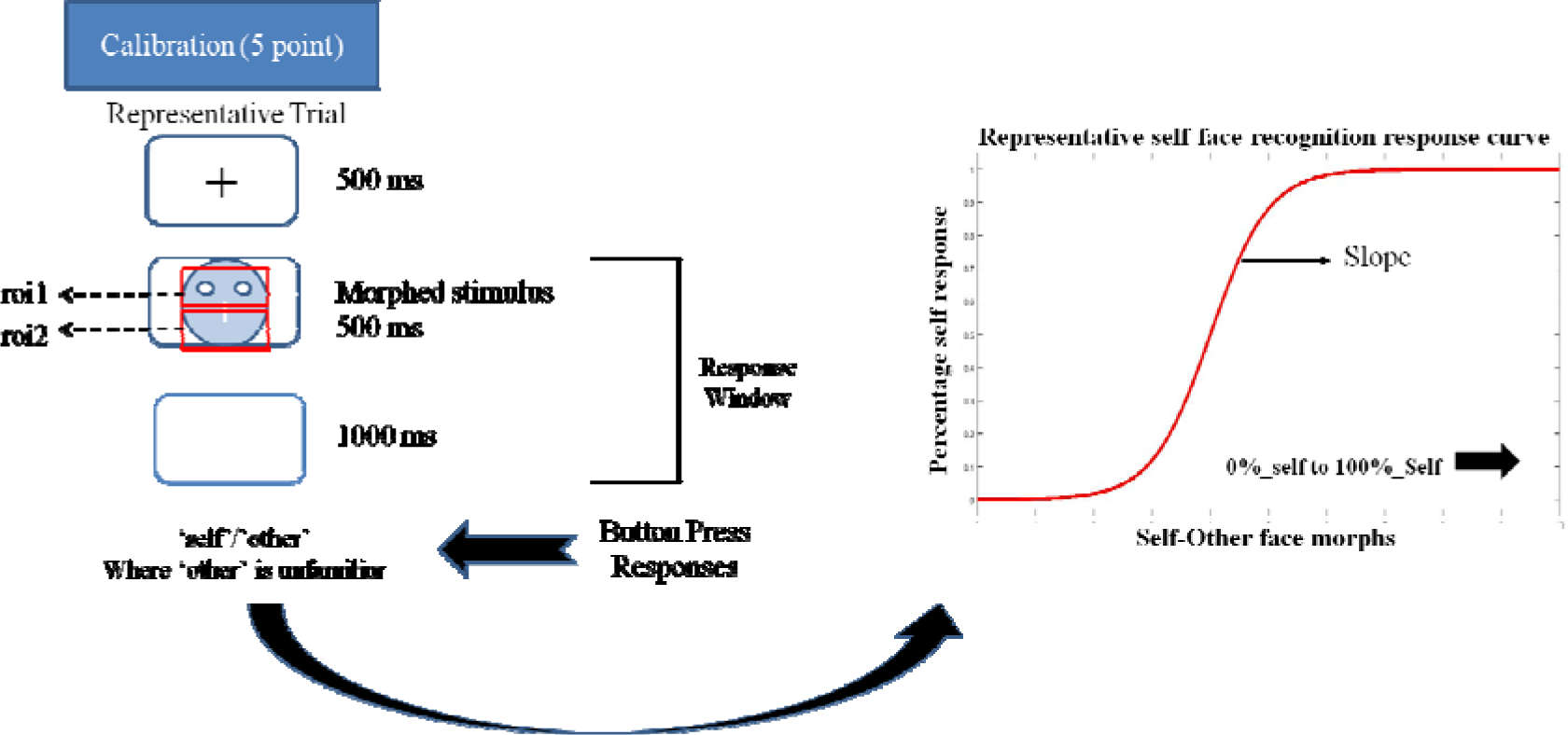
Schematic representation of a trial in the eye tracking task. Participant’s eye movements and gaze pattern were recorded during the 500ms stimulus presentation window and the behavioural response was recorded in the 500 +1000ms window. Participants pressed the ‘a’ key for identifying a face as ‘self’ and ‘l’ key for identifying a face as ‘other’ in the self left-hand response. These key responses were reversed for the self-right hand response. The schema also shows a representative self-face recognition response curve calculated from the ‘self’/other’ face recognition data.

It is predicted that the slope of the self – recognition response curve will be positively associated with the proportion of gaze duration to upper parts of the morphed faces identified as ‘self’ in comparison to faces identified as other. Since eyes provide the richest source of information for identification of a face (Laughery, Alexander et al. 1971, Janik, Wellens et al. 1978, Emery 2000, Schyns, Bonnar et al. 2002, Henderson, Williams et al. 2005, Itier, Alain et al. 2007, Luria and Strauss 2013), it is predicted that individuals who spend more time extracting information from the eyes for faces identified as ‘self’ will have a more distinct self-face representation as evidenced by a steeper self-recognition response slope. They are expected to perform less facial feature sampling of the mouth or other surrounding regions. This processing strategy is not predicted for the unfamiliar other face as there is no previous exposure to the unfamiliar face that will direct such gaze behaviour.

The third aim is to investigate the association between autistic traits and gaze duration to upper and lower parts of self and other faces. It is predicted that autistic traits will correlate negatively with gaze duration to the upper portion of the face. This negative association between autistic traits and gaze duration to eye-region is predicted to be stronger for faces identified as ‘other’ compared to those identified as ‘self’. One of the theoretical explanations of reduced gaze to the eyeregion in ASD is argued to be a negative and stressful reaction to eye-contact in individuals with ASD(Hutt and Ounsted 1966, Kliemann, Dziobek et al. 2010), a reaction that may not hold true for self-faces.

Alternatively, the social motivation theory of ASD posits that reduced fixation to eye region is driven by reduced reward values associated with social stimuli in ASD (Chevallier, Kohls et al. 2012) with an increased preference for geometrical images compared with social images observed in children with ASD (Pierce, Conant et al. 2011, Pierce, Marinero et al. 2016). If this theory holds true it can be expected that the association between higher autistic traits with reduced gaze to eye regions to be less severe for faces identified as ‘self’. This is predicted because self-face can be argued to be of higher reward value (Devue, Van der Stigchel et al. 2009).

## Methods

### Participant Details

Thirty-three healthy adults (5 males; mean ± SD age = 20.67 ± 3.69 years) were drawn from in and around the University of Reading campus, and received either a small compensation or credit points for their participation. All participants were right-handed and had normal or corrected to normal vision. None of the participants had a current clinical diagnosis of neurological or psychiatric disorder and did not self-report any mental health problems. Ethical approval for the study was obtained from the Department of Psychology Research Ethics Committee of the University of Reading and all methods were carried out in accordance with these guidelines regarding all relevant aspects, such as recruitment, compensation and debriefing of participants, as well as the nature of the experiments and other collected information. All participants provided written informed consent in accordance with the Declaration of Helsinki.

### Stimuli

Stimuli were individually tailored for each participant. Each participant was photographed (Canon PowerShot SX700 HS digital camera) looking directly at the camera and holding a neutral expression. Participants were seated at a distance of 100 cm, under constant artificial lighting and with a white background. One unfamiliar ‘other’ identity for each gender and from the same race and age range was also photographed under similar condition.

Following this, each participant’s photograph was converted to grayscale and external features (hairline, jaw line, and ears) were removed. This photograph was then mounted on an oval frame and cropped to a dimension of 350 × 500 pixels using GIMP(2013). A set of stimuli was created for each participant’s face, by morphing self-face with an ‘unfamiliar faces’ using Sqirlz Morph (Xiberpix, Solihull, UK). The following step sizes were used to create the morphing continuum from 100% to 0% of participant’s face (100, 90, 80, 70, 65, 60, 55, 50, 45, 40, 35, 30, 20, 10, 0). Since the previous data showed that individual differences in self-other face category boundary lie within the morph range of 60 and 30 morph percentages for the self-face recognition task(Chakraborty and Chakrabarti 2015), the morph percentages were at 10% step sizes between 100 and 70 and 30 and 0 and 5% step sizes between 70 and 30.

### Apparatus

Calibration and task presentation was controlled using E-prime 2.2 (Psychology Software Tools, PA, USA) presented with TobiiStudio on a Tobii T60 eye tracker monitor (operating at 60Hz) and resolution of 1024 × 768 pixels. Participants sat in a chair 50 cm from the monitor. They used a keyboard for their responses to the task.

### Eye-tracking Measurements

Before commencing the task, participants underwent a five-point calibration procedure implemented on Tobii Studio. Two regions of interest were pre-positioned over each morphed face for each individual participant. The region of interest 1 (*UPPER ROI*) covered the upper portion of the face including the eyes. The region of interest 2 (*LOWER ROI*) covered the lower portion of the face including the mouth. Gaze within these regions was recorded during this task (see Fig. 1). All ROIs were of the same size.

Next, participants completed a self-face recognition task. Each trial started with fixation cross followed by the stimulus image and then a blank screen (see Fig. 1). Participants were instructed to judge and identify a presented face as either ‘self’ or ‘other’. There were two runs, one for a right handed and the other for a left-handed ‘self’ recognition response. Participants were asked to respond with the right or left hand in a given run. Each run consisted of 15 distinct morphs presented ten times each, resulting in 150 trials per run. Any keyboard response in the 1500 ms window was recorded.

Faster recognition of self compared to other faces has been associated with right hemispheric dominance, i.e. people are slightly quicker to recognise self faces when people respond with their left hand (Keenan et al., 1999). We collected responses from both hands to ensure that the effects of interest were not influenced by similar potential hemispheric dominance effects.

All participants completed the AQ questionnaire online following the completion of the task.

### Data Analysis

Statistical tests were conducted and plots generated using SPSS 21 (IBM SPSS Statistics version 21) and R using ggplot2 package (Wickham 2009).

Slope calculation for self-other recognition: ‘Self’ and ‘other’ responses for both runs were combined for each morph level to generate percentage response curves for self-face recognition response for each participant. The slope of self-recognition for each participant was calculated using a logistic psychometric function fitted for maximum likelihood estimation for Weibull distribution. Depending on the stimulus-related information change across the different morph levels required by an individual participant to shift from the self to other category, the psychometric function gives a steep or shallow slope (see Fig. 1). The steepness of this slope is interpreted as an extent of overlap between the self-face and other face representation. A steeper slope indicates a reduced overlap between the self and other representation. In other words, a steeper slope represents a more distinct self-representation.

AQ score for each participant was calculated using the formula as suggested by the authors (Baron-Cohen, Wheelwright et al. 2001).

*Gaze duration analysis:* Gaze position as well as the regions of interest where gaze was on was recorded using E-prime for each time stamp. Gaze position was determined by averaging the locations of both eyes. In the absence of one eye position during the time stamp, the eye position for the single recorded eye was used. The data were processed using MATLAB. The following criteria were used to identify fixations to be included in the analysis:

1) Three successive time stamps within 35 pixels of the original time stamp. Each time stamp is approximately 16.6ms long, hence for a fixation to be included the eye position needed to be within the region of interest for a minimum of 50ms.
2) If gaze was outside the range for one time stamp (possibly due to blinking) or was not recorded but the following time stamp was inside the range, the fixation was considered legitimate.

Following the gaze position analysis, the average gaze duration to ***UPPER ROI*** was calculated for each participant for all trials that the participant identified as ‘self-face’ (*Average_upper_duration_self)* and ‘other-face’ (*Average_upper_duration_other*). Similarly, the average gaze duration to ***LOWER ROI*** was calculated for each participant for all trial identified as ‘self-face’ *(Average_lower_duration_self)* and ‘other-face’ *(Average_lower_duration_other).*

Next, the proportion of gaze duration to *UPPER ROI* (*Upper_proportion_self*) was calculated for each participant for all trials identified as ‘self-face’ by dividing

Average_upper_duration_self by the sum of Average_upper_duration_self and

Average_lower_duration_self (see Box 1). A similar calculation was done for faces identified as ‘other’. The denominators in both instances were chosen to control for individual differences in total looking time to the different ROIs.

### Data Analysis

*Normality checks and exclusions.* The distribution of all variables was tested before analysis, using Shapiro-Wilkinson’s test of normality. Parametric and non-parametric tests of statistical inference were used accordingly (See Table 1). Influence measures (Cook’s D and leverage) were calculated for each correlation and data points exceeding a cut-off of 4/N were excluded from correlation analysis. Due to the strong directionality of the predictions, 1-tailed statistics are used except for the exploratory analysis between AQ and gaze behaviour.

**Table 1.**
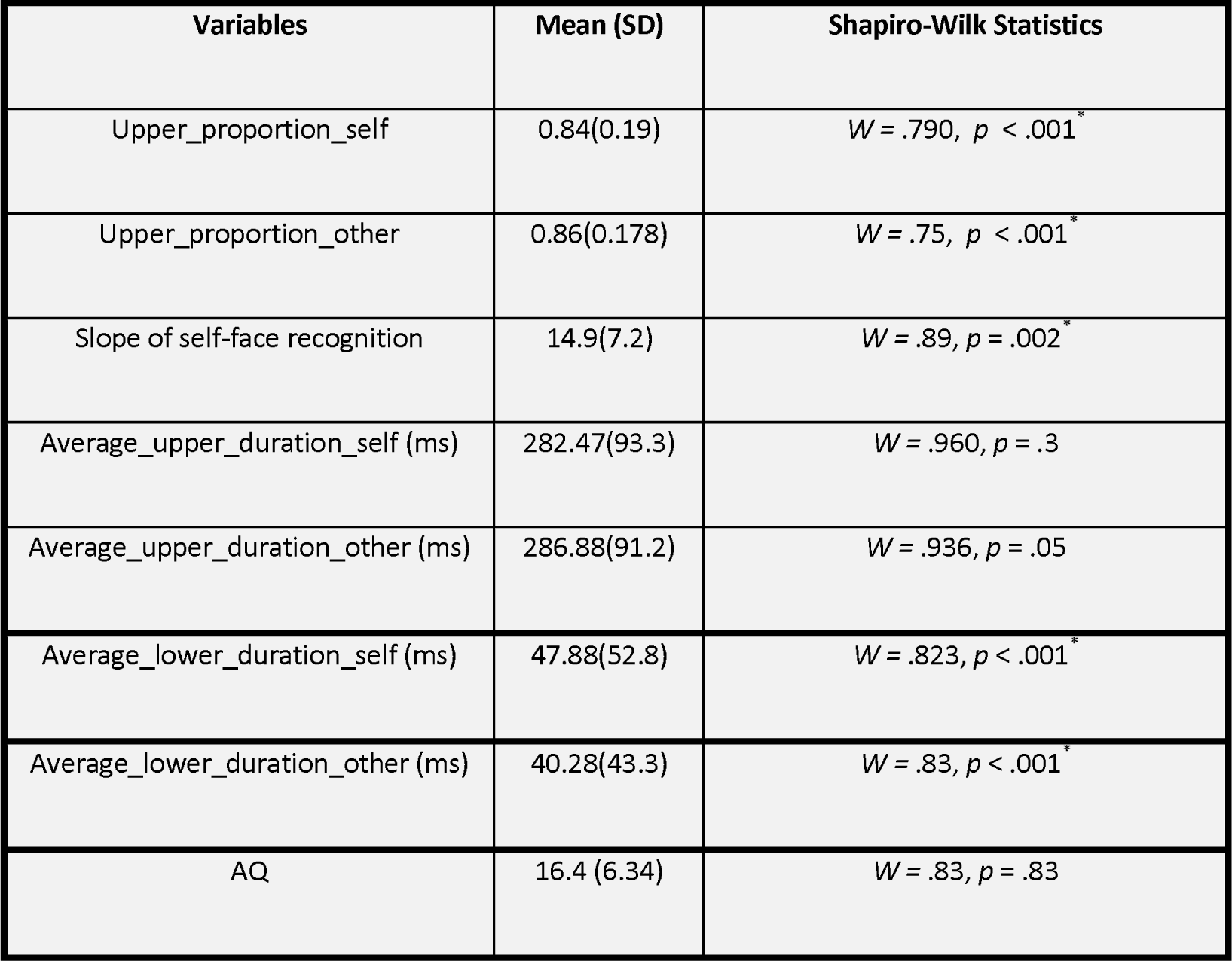
Mean and SD for the computed variables. As shown above, multiple variables violated the assumption of normality. Statistical tests were chosen accordingly. In line with directional nature of the hypothesis, all tests were one-tailed unless specified otherwise.

*Main effects analysis.* To investigate the difference in gaze duration to the upper parts of the face, for faces identified as ‘self’ and ‘other’ a related sample Wilcoxon signed-rank test was computed between Upper_proportion_self and Upper_proportion_other.

*Individual differences analysis.* To investigate individual differences in the association between the slope of the self-recognition curve and gaze duration, Kendall rank correlations were computed between the slope of the self-face recognition response curve and a) the Upper_proportion_self and b) Upper_proportion_other. To investigate individual differences in the association between AQ scores and eye gaze duration, Kendall rank correlations were computed between AQ and a) Upper_proportion_self and b) Upper_proportion_other.

## Results

### Main effects

There was a significantly greater proportion of gaze duration to *UPPER ROI* for morphed faces identified as ‘other’ compared to morphed faces identified as ‘self’ (Wilcoxon Signed-Rank test for Upper_proportion_self & Upper_proportion_other: Z = −2.385, Asymp.Sig = .02, effect sizer = .42). This conversely illustrated that greater proportion of gaze duration to LOWER ROI for morphed faces identified as ‘self’ compared to morphed faces identified as ‘other’.

To investigate if individuals who have a more distinct self-face representation would gaze longer at the upper parts of faces identified as ‘self’ in proportion to faces identified as ‘other’, the ratio of gaze duration for Upper_proportion_self to Upper_proportion_other was chosen as the dependent variable. This ratio was chosen as it provides, for each participant, a quantified response of whether the looking time to the upper parts of faces is higher for faces identified as self vs. those identified as ‘other’. In line with the prediction, a significant positive correlation was observed between the slope of self-face recognition with the ratio of Upper_proportion_self to Upper_proportion_other (Kendall’s tau: *τ* (29) = .23, *p* =.04; see Fig. 2).

**Fig. 2.**
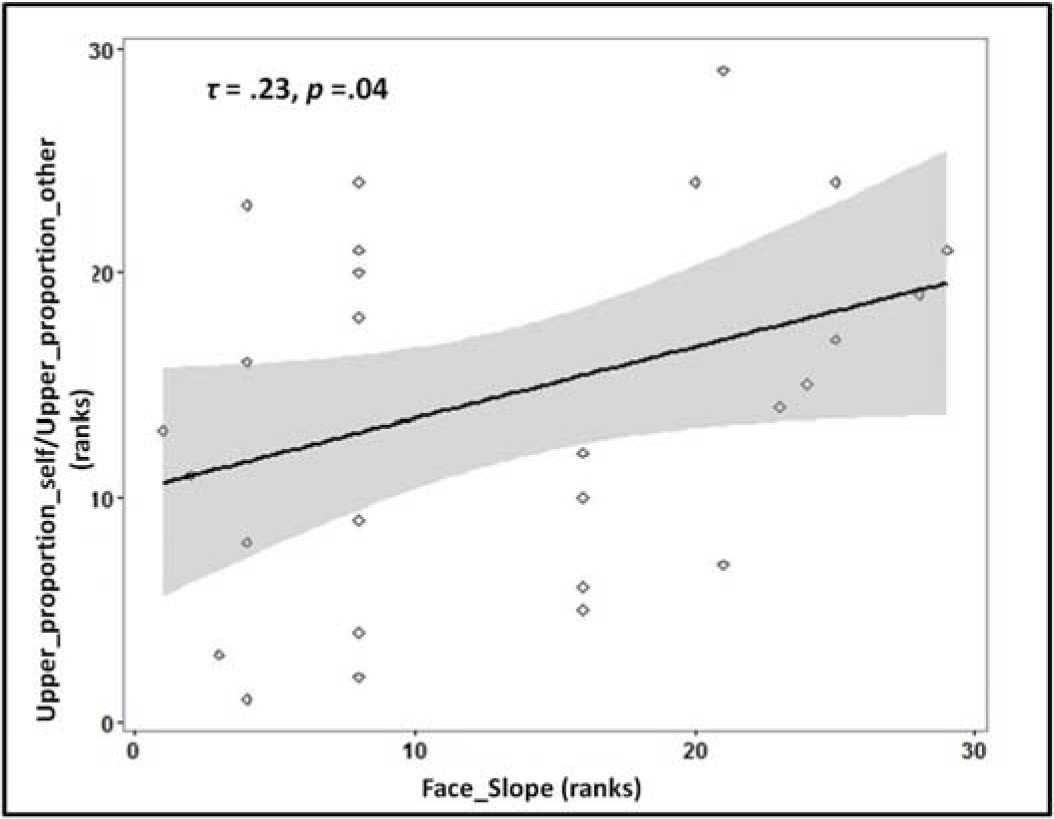
Rank scatterplot representing the positive association between the slope for self-face recognition (x-axis) with the proportion of gaze duration (y-axis) to *UPPER ROI* (ratio_proportion (self:other)) for faces identified as ‘self’ compared to faces identified as ‘other’. The shaded portion represents the 95% confidence region of the slope of the regression line.

No effect of the responding hand (left/right) was noted on the gaze duration to faces identified as ‘self’ or ‘other’ (p>0.05).

### Individual differences

No significant association between autistic traits and proportion of eye gaze duration to *UPPER ROI* for faces identified as ‘self’ (Upper_proportion_self; Kendall’s tau: *τ* (33) = -.008, p = .48) or ‘other’ (Upper_proportion_other; Kendall’s tau: *τ* (33) = .02, *p* = .45).

Following previous findings of reduced overall looking time to social stimuli (like faces) in individuals with ASD, an exploratory analysis was carried out to investigate if the overall looking time to faces (adding the gaze duration for both regions of interest) was associated with autistic traits. The total gaze duration (for each participant) was calculated for faces identified as ‘self’ and as ‘other’. A significant negative correlation was observed between autistic traits and total looking time for faces identified as ‘self’ (Kendall’s tau: *τ* (33) = -.305, *p* =.01) as well as faces identified as ‘other’ (Kendall’s tau: *τ* (33)= -.286, *p* =.02; see Fig. 3).

**Fig. 3.**
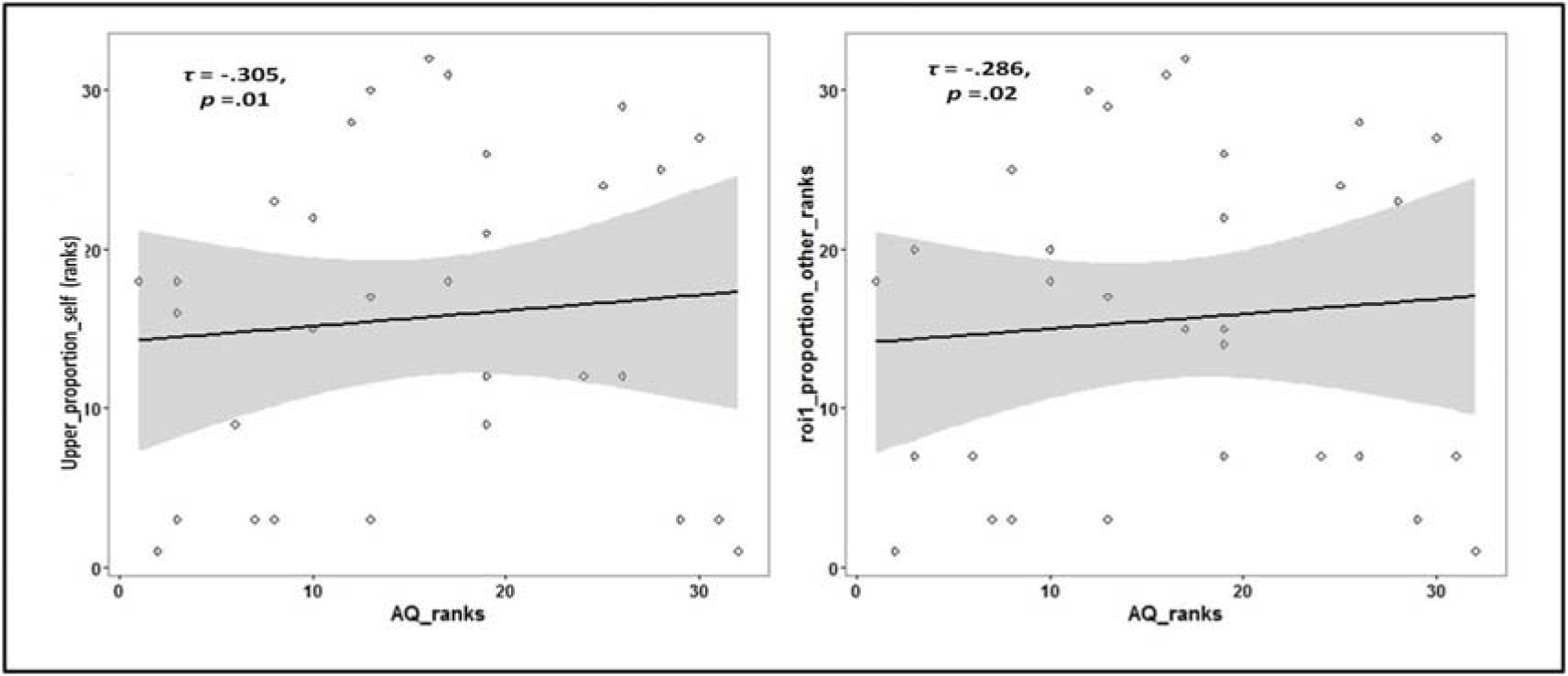
Rank scatterplots representing the negative association between the AQ scores with the total gaze duration for faces identified as a) ‘self’ and for faces identified as b) ‘other’. The shaded portion represents the 95% confidence region of the slope of the regression line

Since the denominators (total looking time) in the ratios (Upper_proportion_self & Upper_proportion_other) were negatively correlated with autistic traits, this might have influenced the overall relation between the proportion of gaze duration and autistic traits. Hence further correlation analyses were performed between AQ and the average gaze duration for both ROIs and both face categories. A negative association trend was observed between autistic traits and average gaze duration to *UPPER ROI* for faces identified as ‘self’ and for faces identified as ‘other’ (Kendall’s tau: *τ* (33) =-.197, *p*=.06). No such trend was observed between autistic traits and average gaze duration to *LOWER ROI* for both faces identified as ‘self’ and faces identified as ‘other’ (Kendall’s tau: *τ* (33) = -.041, *p* = .4).

In line with previous results (Chakraborty and Chakrabarti 2015), no significant association was observed between the self-face recognition slope and autistic traits (Kendall’s tau = *τ* (33) = -.120, *p* = 2).

## Discussion

This study tested differences in gaze pattern for faces identified as ‘self’ and ‘other’ from a series of self-other face morphs. The study also investigated the association of these gaze patterns with the behavioural representation of self-faces and autistic traits. In line with the prediction, we found a significant difference in gaze duration to upper vs. lower regions of a face between faces that were identified as ‘self’ and those identified as ‘other’. We also found that individuals with a more distinct self-face representation looked longer at the upper region of faces identified as self vs. those identified as other. Contrary to our predictions, no significant association was observed between autistic traits and the proportion of gaze duration to upper parts of morphed faces for faces identified as ‘self’ or for faces identified as ‘other’. However, a negative association between autistic traits and total gaze duration to both faces identified as ‘self’ and as ‘other’ was noted. The results are discussed in details in the following paragraphs.

### Increased facial feature sampling for faces labelled as ‘self’ compared to those labelled as ‘other’

Self-face has high relational salience (Breédart, Delchambre et al. 2006) to the individual and may possess high subjective reward value (Devue, Van der Stigchel et al. 2009). However, the visual processing strategies employed in recognising the highly salient and familiar self-face is relatively unknown. The current study found that faces identified as ‘self’ had increased gaze duration to lower parts of the face compared to faces identified as ‘other’. However, the gaze duration to the upper region of the face was not significantly different between the two identities. In line with this result, a previous eye tracking study reported increased feature sampling for familiar faces across different regions of the face (Van Belle, Ramon et al. 2010). The increased sampling of features for faces labelled as ‘self’ is in line with the similar observation reported for familiar faces. This observation is arguably driven by the salience and reward value of the self face as a stimulus. Previous work has also found individuals maintain sustained attention to the self-face which can interfere with concomitant task performance(Devue, Van der Stigchel et al. 2009).

### Greater upper region sampling of faces labelled as ‘self’ associated with a more distinct self-face representation

The slope of the psychometric function for self-recognition was positively associated with the ratio of gaze proportion to the upper region for faces identified as ‘self’ to those identified as ‘other’. This finding suggests that individuals with a more distinct self-face representation spent a greater proportion of time looking at the upper part of faces (including the eye region) for faces labelled as ‘self’. Due to the correlational nature of the study, it is not possible to infer directionality of this association. This finding opens up questions about the stability of self-face representation, and the impact of task manipulations on it. If self-face representation is influenced by task conditions, a future experiment could explicitly ask participants to look at the upper vs. lower regions of the face, to test if that alters the slope of the self-face representation.

The current study did not compare self-face with familiar other faces. Depending on the exposure level, a familiar face may also be of high salience and well represented mentally. Follow-up research should test if distinct behavioural representations of familiar other faces are associated with increased gaze duration to upper parts of these faces.

### Gaze duration to faces is reduced with higher autistic traits

Reduced gaze duration to both self and other faces was noted in individuals with high autistic traits. This observation replicates several previous reports where individuals with ASD have been shown to demonstrate reduced gaze duration toward faces(Pelphrey, Sasson et al. 2002, Dalton, Nacewicz et al. 2005). However, no significant difference in this association was noted for self vs. other faces. This observation suggests that a) individuals are performing at ceiling due to the relative ease of the task, thus masking any potential difference between the processing of self vs. other faces, or b) the differences in gaze processing strategies between self and other faces are orthogonal to the dimension of autistic traits. A previous study investigating self, familiar, and unfamiliar face viewing using eye tracking did not observe any difference in gaze fixation pattern to different categories of faces between control and ASD children(Gillespie-Smith, Doherty-Sneddon et al. 2014). However, the association was observed between socio-communicative abilities and gaze patterns to self and unfamiliar faces. The current study is consistent with this earlier report.

One limitation of the current study is that it did not include non-self familiar faces as a stimulus category. In order to directly compare the self-face related results from this study with previous eye-tracking studies investigating familiar face recognition, future experiments should include both of these conditions in the same task. A second limitation of this task was not optimising it for measuring response times. Participants were not given any instruction on how quickly to respond, which might have led to different strategies employed by different participants. Finally, the scope the current study is also limited in terms of the trait measures that it investigates. While autistic traits are definitely of interest to self-face processing, they are by no means the only dimensions that can be theoretically linked to potential differences in these processes. Future research should examine other traits that could relate to individual differences in self face representation and associated gaze behaviour to self faces. For example, do individuals who exhibit a more distinct self-face representation exhibit preoccupation with their body image?

## Conclusion

This study shows that the visual processing of faces identified as ‘self’ is different from those identified as an unfamiliar other. These differences in visual processing are associated with individual differences in self-face representation. Individuals with a ‘more distinct’ self-face representation spent a greater proportion of time looking at the upper regions of faces identified as self, compared to those identified as other. The results from this study support the idea that self-faces might be processed similarly to other familiar faces, and self-specificity effects may come into play at higher order relay regions in the brain(Kita, Gunji et al. 2010).

This study also shows that higher autistic traits do not specifically influence gaze responses to self-face but reduce looking time to faces (self and other) in general (see Fig. 4 for a summary of results). Consistent with a previous report (Chakraborty and Chakrabarti 2015) in a similar neurotypical population, this study found that psychometric properties of physical self-representation, appears to be uninfluenced by autistic traits. This observation lends support to the domain-specific influence of autistic traits on self-representations (Williams 2010). However, physical self-representation needs to be formally tested in a clinically diagnosed ASD population using similar approaches to test the generalisability of the current results to the extreme high end of the spectrum of autistic traits.

**Fig. 4.**
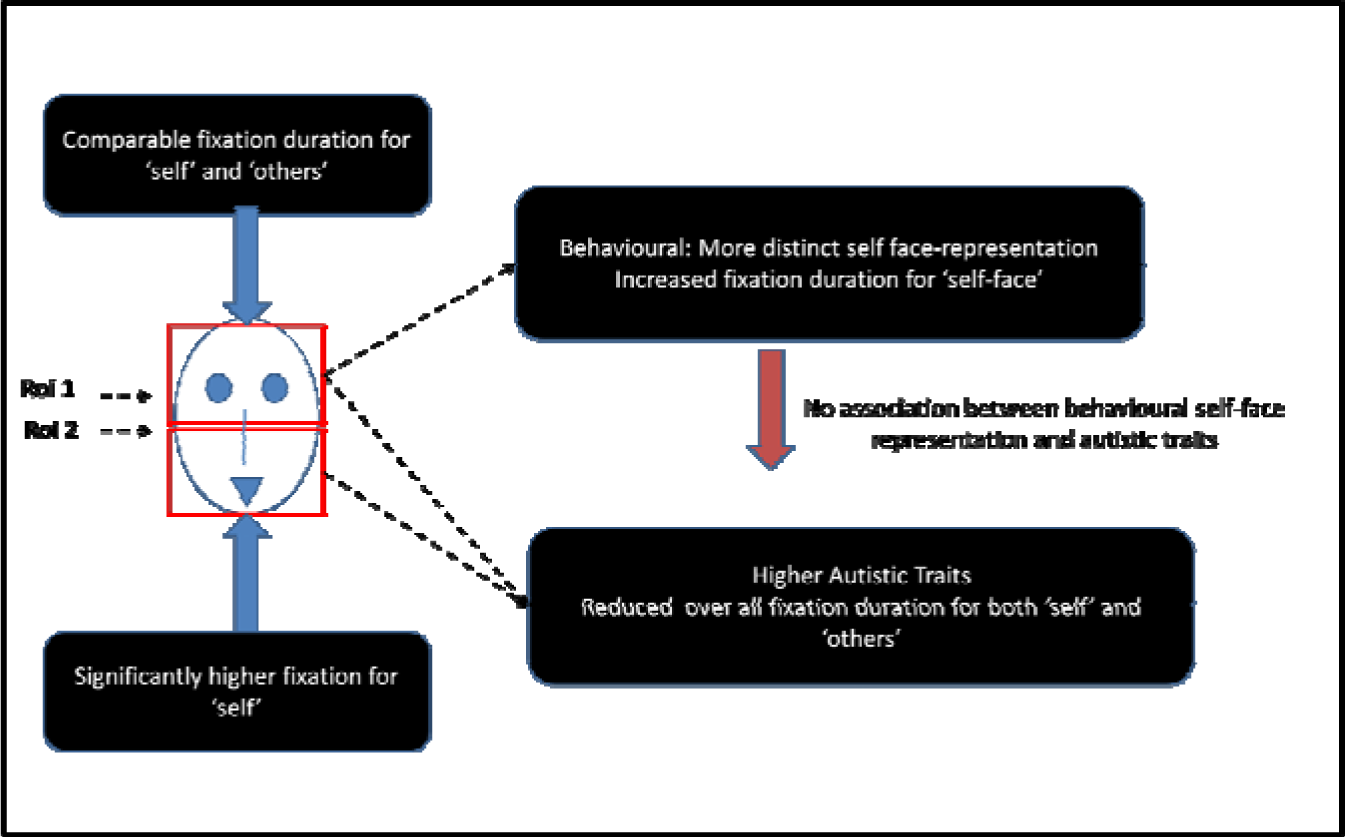
Schematic representation of the main findings from the study

## Acknowledgements

This work was supported by Felix Scholarship Trust. BC was supported by the Medical Research Council UK. The authors wish to thank Anthony T Haffey and Christopher P. Taylor for help with data analysis.

## Author Contributions

Both authors developed the study concept. The study design, data collection, analysis, interpretation and draft of manuscript were performed by A.C. under supervision and critical revisions of B.C.

Both authors approved the final version of the manuscript for submission.

## Additional Information

Competing financial interests: The authors declare no competing financial interests.

## References

Amsterdam, B. (1972). “Mirror self-image reactions before age two.” Developmental psychobiology 5(4): 297–305, doi: 10.1002/dev.420050403

Baron-Cohen, S., S. Wheelwright, R. Skinner, J. Martin and E. Clubley (2001). “The autism-spectrum quotient (AQ): Evidence from asperger syndrome/high-functioning autism, malesand females, scientists and mathematicians.” Journal of autism and developmental disorders 31(1): 5–17, doi: 10.1023/A:1005653411471

Breédart, S., M. Delchambre and S. Laureys (2006). “Short article one’s own face is hard to ignore.” The Quarterly Journal of Experimental Psychology 59(1): 46–52, doi: 10.1080/17470210500343678

Bird, G. and Viding, E (2014). “The self to other model of empathy: providing a new framework for understanding empathy impairments in psychopathy, autism, and alexithymia.” Neuroscience Biobehaviour Review 47(11): 520–532, doi: 10.1016/j.neubiorev.2014.09.021

Chakraborty, A. and B. Chakrabarti (2015). “Is it me? Self-recognition bias across sensory modalities and its relationship to autistic traits.” Molecular autism 6(1): 1, doi: 10.1186/s13229-015-0016-1

Chevallier, C., G. Kohls, V. Troiani, E. S. Brodkin and R. T. Schultz (2012). “The social motivation theory of autism.” Trends in cognitive sciences 16(4): 231–239, doi: 10.1016/j.tics.2012.02.007

Dalton, K. M., B. M. Nacewicz, T. Johnstone, H. S. Schaefer, M. A. Gernsbacher, H. Goldsmith, A. L. Alexander and R. J. Davidson (2005). “Gaze fixation and the neural circuitry of face processing in autism.” Nature neuroscience 8(4): 519–526, doi: 10.1038/nn1421

Devue, C., S. Van der Stigchel, S. Breédart and J. Theeuwes (2009). “You do not find your own face faster; you just look at it longer.” Cognition 111(1): 114–122, doi: 10.1016/j.cognition.2009.01.003

Emery, N. J. (2000). “The eyes have it: the neuroethology, function and evolution of social gaze.” Neuroscience & Biobehavioral Reviews 24(6): 581–604, doi: 10.1016/S0149-76340000025-7

Gallup, G. G. (1970). “Chimpanzees: self-recognition.” Science 167(3914): 86–87, doi: 10.1126/science.167.3914.86

Gillespie-Smith, K., G. Doherty-Sneddon, P. J. Hancock and D. M. Riby (2014). “That looks familiar: attention allocation to familiar and unfamiliar faces in children with autism spectrum disorder.” Cognitive neuropsychiatry 19(6): 554–569, doi: 10.1080/13546805.2014.943365

Gillihan, S. J. and M. J. Farah (2005). “Is self special? A critical review of evidence from experimental psychology and cognitive neuroscience.” Psychological bulletin 131(1): 76, doi: 10.1037/0033-2909.131.1.76

“GNU Image Manipulation Program.” from http://www.gimp.org/.

Gray, H. M., N. Ambady, W. T. Lowenthal and P. Deldin (2004). “P300 as an index of attention to self-relevant stimuli.” Journal of Experimental Social Psychology 40(2): 216–224, doi: 10.1016/S0022-1031 (03)00092-1

Heinisch, C., H. R. Dinse, M. Tegenthoff, G. Juckel and M. Bruüne (2011). “An rTMS study into self-face recognition using video-morphing technique.” Social cognitive and affective neuroscience 6(4): 442–449, doi: 10.1093/scan/nsq062

Heisz, J. J. and D. I. Shore (2008). “More efficient scanning for familiar faces.” Journal of Vision 8(1): 9–9, doi: 10.1167/8.1.9

Henderson, J. M., C. C. Williams and R. J. Falk (2005). “Eye movements are functional during face learning.” Memory & Cognition 33(1): 98–106, doi: 10.3758/BF03195300

Hungr, C. J. and A. R. Hunt (2012). “Physical self-similarity enhances the gaze-cueing effect.” The Quarterly Journal of Experimental Psychology 65(7): 1250–1259, doi: 10.1080/17470218.2012.690769.

Hutt, C. and C. Ounsted (1966). “The biological significance of gaze aversion with particular reference to the syndrome of infantile autism.” Behavioral science 11(5): 346–356, doi: 10.1002/bs.3830110504

Itier, R. J., C. Alain, K. Sedore and A. R. McIntosh (2007). “Early face processing specificity: It’s in the eyes!” Journal of Cognitive Neuroscience 19(11): 1815–1826, doi:10.1162/jocn.2007.19.11.1815

James, W. (1890). “The principles of psychology (Vol. I).” New York: Holt.

Janik, S. W., A. R. Wellens, M. L. Goldberg and L. F. Dell’Osso (1978). “Eyes as the center of focus in the visual examination of human faces.” Perceptual and motor skills 47(3): 857–858, doi: 10.2466/pms.1978.47.3.857

Keenan, J. P., G. Ganis, S. Freund and A. Pascual-Leone (2000). “Self-face identification is increased with left hand responses.” Laterality: Asymmetries of Body, Brain and Cognition 5(3): 259–268, doi: 10.1080/713754382

Keenan, J. P., B. McCutcheon, S. Freund, G. G. Gallup, G. Sanders and A. Pascual-Leone (1999). “Left hand advantage in a self-face recognition task.” Neuropsychologia 37(12): 1421–1425 doi: 10.1016/S0028-3932(99)00025-1

Keenan, J. P., M. A. Wheeler and M. Ewers (2003). “The neural correlates of self-awareness and self-recognition.” The self in neuroscience and psychiatry: 166–179, doi: 10.1016/j. concog.2010.09.007

Kircher, T. T., M. Brammer, E. Bullmore, A. Simmons, M. Bartels and A. S. David (2002). “The neural correlates of intentional and incidental self processing.” Neuropsychologia 40(6): 683–692, doi: 10.1016/S0028-3932(01)00138-5

Kircher, T. T., C. Senior, M. L. Phillips, S. Rabe-Hesketh, P. J. Benson, E. T. Bullmore, M. Brammer, A. Simmons, M. Bartels and A. S. David (2001). “Recognizing one’s own face.” Cognition 78(1): B1–B15, doi: 10.1016/S0010-0277(00)00104-9

Kita, Y., A. Gunji, K. Sakihara, M. Inagaki, M. Kaga, E. Nakagawa and T. Hosokawa (2010). “Scanning strategies do not modulate face identification: Eye-tracking and near-infrared spectroscopy study.” PloS one 5(6): e11050, doi: 10.1371/journal.pone.0011050

Kliemann, D., I. Dziobek, A. Hatri, R. Steimke and H. R. Heekeren (2010). “Atypical reflexive gaze patterns on emotional faces in autism spectrum disorders.” The Journal of Neuroscience 30(37): 12281–12287, doi: 10.1523/JNEUR0SCI.0688-10.2010

Robinson, E. B., Koenen, K. C., McCormick, M. C., Munir, K., Hallett, V., Happeé, F., … & Ronald, A. (2011). Evidence that autistic traits show the same etiology in the general population and at the quantitative extremes (5%, 2.5%, and 1%). Archives of general psychiatry, 68(11), 1113–1121, doi: 10.1001/archgenpsychiatry.2011.119

